# Preclinical stage abundance and nuclear antigen reactivity of fecal Immunoglobulin A (IgA) varies among males and females of lupus-prone mouse models

**DOI:** 10.1101/2022.01.26.477917

**Authors:** Radhika Gudi, Soumyabrata Roy, Wei Sun, Chenthamarakshan Vasu

**Author notes:** Address Correspondence: Chenthamarakshan Vasu, Microbiology and Immunology, Medical University of South Carolina, 173 Ashley Avenue, BSB214B, Charleston, SC-29425.

## Abstract

Systemic lupus erythematosus (SLE) is characterized by the production of pathogenic autoantibodies with nuclear antigen (nAg) specificity. Using (SWRxNZB)F1 (SNF1) mice, we showed higher levels of IgA production in the intestine and the nAg reactivity of fecal IgA under lupus susceptibility. Here, we determined if the fecal IgA abundance and nAg reactivity are higher in, different among, various lupus-prone preclinical models (MRL/lpr, NZBxNZW-F1, SNF1, NZM2410 and NZM2328). We also determined if the fecal IgA nAg reactivity at pre-seropositive ages correlates with the eventual serum autoantibody levels in males and females of these mouse models. We show that age dependent increase in the abundance and nAg reactivity of fecal IgA can vary among different lupus-prone mouse models. Importantly, fecal IgA in these mice show significant levels of nAg reactivity, starting as early as at juvenile age. Furthermore, the pre-seropositive stage nAg reactivity of fecal IgA in most lupus-prone strains correlates well with that of eventual, seropositive stage systemic autoantibody levels. Gender differences in serum autoantibody levels were preceded by similar differences in the fecal IgA abundance and nAg reactivity. These observations suggest that fecal IgA features, nAg reactivity particularly, could serve as a biomarker for early prediction of the eventual systemic autoimmunity in lupus-prone subjects.

## Introduction

Systemic Lupus Erythematosus (SLE) is a chronic autoimmune disease which involves various genetic, epigenetic, hormonal, environmental and immune regulatory factors [1–4]. SLE affects mostly women (about 9:1 over men) and the clinical features are unpredictable with periods of remission and flares[4]. SLE arises when abnormally functioning B lymphocytes produce autoreactive antibodies to DNA and various nuclear proteins. Hence, large amounts of circulating autoantibodies as well as immune complex deposition in the kidney causing glomerulonephritis are the key features of SLE [2]. Recent studies including ours, have shown that gut microbiota composition influences the rate of disease progression and the overall disease outcome [5–12]. Using a slow progressing lupus-prone mouse model, (SWRxNZB)F1 (SNF1) strain, we showed a potential contribution of gut microbiota and pro-inflammatory immune response initiated in the gut mucosa in triggering the disease associated gender bias in SLE [12, 13]. We also reported that the preclinical stage IgA production in the gut, and the abundance and nuclear antigen (nAg) reactivity of fecal IgA are correlative of the onset of anti-nAg seropositivity and the eventual proteinuria, and gender bias of the disease incidence in SNF1 mice[14].

IgA produced in the gut plays an important role in fighting microbial infection and maintaining a healthy gut microbiota[15–17]. Higher levels of total IgA in stool samples of SLE patients compared to that of healthy controls have been reported [5]. Importantly, anti-DNA antibodies of IgA class are found in the serum of patients with SLE [18–23]. These reports along with our studies [13, 14, 24] showing pro-inflammatory immune phenotype and higher plasma cell frequency in the intestine, and higher abundance and nAg reactivity of fecal IgA in female SNF1 mice at preclinical stages suggested that the degree of IgA secretion in the gut lumen could be indicative of systemic autoimmune progression. However, whether these features are unique to SNF1 mice or ubiquitous to other preclinical models of lupus [such as MRL/lpr, NZM2328, NZM2410, (NZBxNZW-F1) NZB/WF1] is not known. These mouse strains including SNF1 develop lupus-like disease spontaneously and present varying degrees of gender bias and/or timing of disease onset[13, 25–30]. In the present study, we investigated the pre-clinical stage fecal IgA features such as abundance and nAg reactivity of aforementioned mouse strains. We also examined if fecal IgA features at younger ages show a correlation with the eventual nAg reactivity of serum IgG in males and females. We found that fecal IgA in MRL/lpr, NZM2328, NZM2410, NZB/WF1 mice, similar to SNF1 mice[14], shows varying degrees of age dependent increase in the abundance and nAg reactivity. Fecal IgA in these mice showed considerable levels of nAg reactivity starting at as early as juvenile age. Further, the nAg reactivity of fecal IgA in most of these mouse strains at pre-seropositive stage showed a trend similar to that of eventual circulating nAg reactive IgG levels. These observations suggest that fecal IgA features, nAg reactivity particularly, could serve as a biomarker for early prediction of eventual systemic autoimmunity in lupus-prone subjects at pre-seropositive and pre-clinical stages.

## Materials and methods

### Mice

SWR/J (SWR), NZB/BlNJ (NZB), NZW/LacJ (NZW) and C57BL/6J (B6) mice were purchased from the Jackson Laboratory (Bar Harbor, Maine) and bred under specific pathogen free (SPF) conditions at the animal facilities of Medical University of South Carolina (MUSC). (SWRxNZB)F1 (SNF1) hybrids were generated at the SPF facility of MUSC by crossing SWR females with NZB males. NZB females and NZW males were crossed to generate (NZBxNZW)F1 (NZBWF1) mice. MRL-lpr and NZM2140 breeders were provided by Dr. Gary Gilkeson (MUSC).

NZM2328 breeders were provided by Dr. Shu Man Fu (University of Virginia). Urine and tail vein blood samples were collected from individual mice at different time-points to detect proteinuria and serum autoantibody levels, respectively. All experimental protocols were approved by the Institutional Animal Care and Use Committee (IACUC) of MUSC. All methods in live animals were carried out in accordance with relevant guidelines and regulations of this committee and performed according to National Institutes of Health (NIH) Guidelines on Humane Care and Use of Laboratory Animals.

### ELISA

For determining antibody levels in fecal samples, extracts of fecal pellets collected from individual mice were used. Weighed fecal pellets were suspended in proportionate volume of PBS (100 mg feces/ml PBS; w/v) by breaking the pellet using a pipette tip and high-speed vortex, and continuous shaking at 800 rpm overnight at 4°C. Suspensions were centrifuged at 14,000 rpm for 15min and the top 2/3 of the supernatants were diluted optimally for determining total IgA concentration. Total IgA levels were determined by employing in-house quantitative sandwich ELISA. For SNF1, NZB/WF1, NZM2410 and B6 mice, 1:200 dilution was employed as the optimum dilution for determining total IgA concentration. Optimum dilutions of 1:500 and 1:2000 respectively were employed for NZM2328 and MRL/lpr mice. For this ELISA, purified anti-mouse IgA monoclonal antibody (0.1 μg/well) coated wells were incubated with diluted samples for 2h, incubated with biotin-linked polyclonal anti-mouse IgA antibody for 1h, and finally with streptavidin-HRP for 30min, before developing the reaction using TMB substrate. Purified mouse IgA (Southern biotech) was used in each plate for generating the standard curve.

Antibody titers against nAgs [(double stranded DNA (dsDNA) and nucleohistone (NH)] in mouse sera and fecal extracts were assessed by ELISA as described in our recent reports with minor modifications [9, 12–14]. Briefly, 0.5μg/well of nucleohistone (Sigma-Aldrich) in carbonate buffer or 0.5μg/well dsDNA from calf thymus (Sigma-Aldrich) in 50% DNA coating solution (Thermo) + 50% PBS was coated as nAg onto ELISA plate wells. All sample and antibody reagent dilutions were made in 1% BSA. Serial dilutions of the fecal extracts (starting at 1:20 dilution in 1% BSA) or serum samples (starting at 1:1000 dilution in 1% BSA) were made, IgA or IgG antibodies against these antigens were detected using biotin-conjugated respective anti-mouse antibodies, followed by incubation with streptavidin-HRP (Sigma-Aldrich, Invitrogen and Southern Biotech), and the reaction was detected using TMB substrate (BD biosciences). Known positive and negative control samples identified from the initial screening using antigen-coated and non-coated plates, were used in all plates to validate the results for determining the reliability of the titer values. Highest dilution of the sample that produced an OD value of ≥0.05 above the background value was considered as the nAg reactive titer of that sample. Baseline titer value of 10 was assigned to samples that showed OD values of <0.05 at 1:20 dilution. IgA concentrations (μg/gram feces) and nAg reactive titers (titer/per gram feces) were calculated by multiplying all values by 10.

### Proteinuria

Urine samples were tested at regular intervals, as early as at 12 weeks of age, to detect proteinuria and confirm the previously reported trends in disease incidence in different strains of lupus-prone mice from our facility. Protein level in the urine was determined by Bradford assay (BioRad) against bovine serum albumin standards as described before[9, 13, 14]. Mice that showed proteinuria (>5 mg/ml) were considered to have severe nephritis or clinical stage disease.

### Statistical analysis

GraphPad Prism or Microsoft excel was used to calculate the statistical significance. Mann-Whitney test was employed to calculate *p*-values when two means were compared. Proteinuria positive and negative male and female mice within each strain were compared statistically by Chi-square test. Pearson’s correlation coefficient/bivariate correlation approach was employed for measuring linear correlation between two variables. A *P* value ≤0.05 was considered statistically significant.

## Results

### Fecal IgA abundance varies among lupus-prone mouse strains

Our recent report [14] showed that the abundance and nAg reactivity of fecal IgA are higher in lupus-prone SNF1 mice and these features present gender bias at as early as juvenile and pre-seropositive ages. In this study, we examined if such IgA features (higher abundance and nAg reactivity) are detectable in other major strains of lupus-prone (MRL/lpr, NZM2328, NZM2410 and NZB/WF1) mice as well. Of note, long term-monitoring of the mice used in this study for proteinuria showed previously reported[12, 13, 26, 27, 30, 31] trends in clinical disease onset (**Supplemental Fig. 1**).

Fresh fecal samples from lupus susceptible, individual male and female MRL/lpr, NZB/WF1, SNF1, NZM2410 and NZM2328 mice at different ages (4, 8 and 12 weeks of age) were collected, cryopreserved, and used for preparing fecal extracts and tested for total IgA levels. Fecal samples from male and female B6 mice (lupus non-susceptible controls) were tested similarly. As observed in **Supplemental Fig. 2**, total IgA antibody levels were significantly higher in all lupus-prone mouse strains as early as juvenile age (4 weeks) compared to age matched B6 mice. As observed in **Fig. 1**, all lupus-prone mouse strains, except NZB/WF1, showed some degree of age-dependent increase in fecal IgA levels at adult ages compared to their juvenile age levels. Among these lupus-prone mice, while the NZM2410 and SNF1 mice showed the lowest amounts of fecal IgA at juvenile and adult ages respectively, MRL/lpr mice showed the highest abundance of fecal IgA at all ages. Importantly, both NZB/WF1 and SNF1 females showed significantly higher levels of fecal IgA compared to that of their male counterparts during adult ages. MRL/lpr, NZM2410 and NZM2328 mice, although showed similar overall trends, gender specific differences in fecal IgA levels were not statistically significant in them. These observations show that the overall amounts of IgA secreted in the fecal samples are significantly higher in lupus-prone mouse models and significant gender bias in the fecal IgA levels is detectable in NZB/WF1 and SNF1 strains.

**Figure 1:**
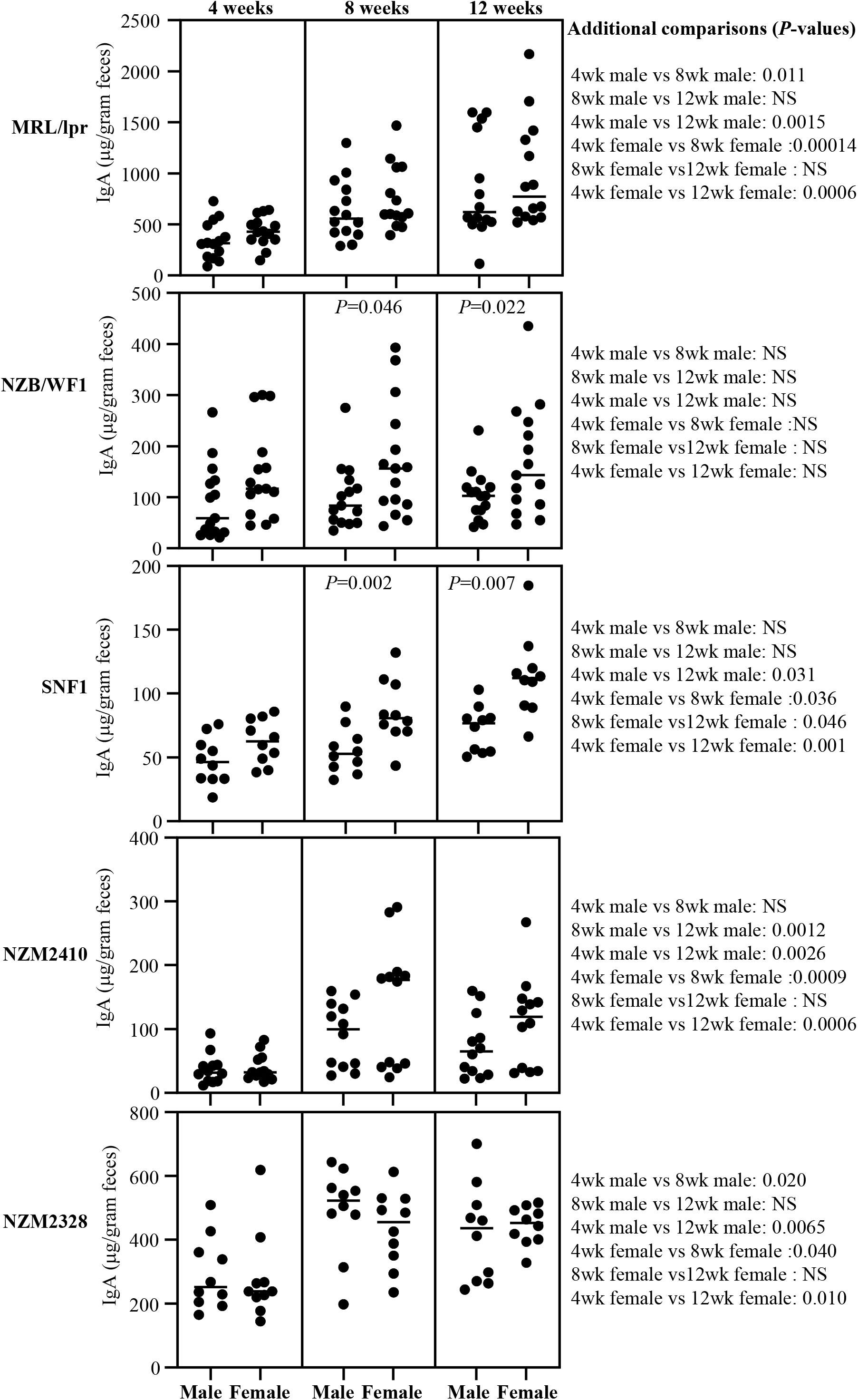
Abundance of IgA in the feces of lupus-prone mouse strains. Fecal samples were collected from individual mice of indicated strains at 4, 8 and 12 weeks of age and subjected to ELISA to determine the IgA concentrations as detailed under materials and method section. IgA concentrations per gram feces are shown. n= 14 males and 14 females for MRL/lpr, 15 males and 15 females for NZB/WF1, 10 males and 10 females for SNF1, 12 males and 12 females for NZM2410, and 10 males and 10 females for NZM2328 strains. *P*-value (by Mann-Whitney test) of male vs female comparisons are shown within the graph. *P*-values of other comparisons are shown on the right.

### Lupus-prone mice show fecal IgA with nAg reactivity at as early as juvenile age

dsDNA and NH reactivities of fecal IgA from various strains of lupus-prone mice as well as B6 control mice were examined. **Supplemental Fig. 3** shows that nAg reactivity (anti-dsDNA and anti-NH) titers of fecal IgA in all lupus-prone mouse strains, in females particularly, are higher at juvenile age (4 weeks), as compared to non-susceptible B6 mice, and this difference is amplified at adult age (12 weeks). **Figs. 2 and 3** show that nAg (dsDNA and NH) reactive fecal IgA titers are profoundly higher in MRL/lpr mice compared to that of other lupus-prone strains. Importantly, all lupus-prone mice showed an overall age-dependent increase in nAg reactivity of fecal IgA at adult ages, as compared to their juvenile age nAg reactive IgA levels. Importantly, dsDNA and NH reactivities of fecal IgA were profoundly higher in female NZB/WF1 and SNF1 mice compared to their male counterparts, particularly at adult ages. However, no statistically significant differences in nAg reactive fecal IgA levels were observed between males and females of MRL/lpr, NZM2410 and NZM2328 mice. Overall, these results show that nAg reactivity of fecal IgA is higher in all lupus-prone mice compared to B6 mice and significant gender differences in nAg reactivities of fecal IgA can be detected in NZB/WF1 and SNF1 strains.

**Figure 2:**
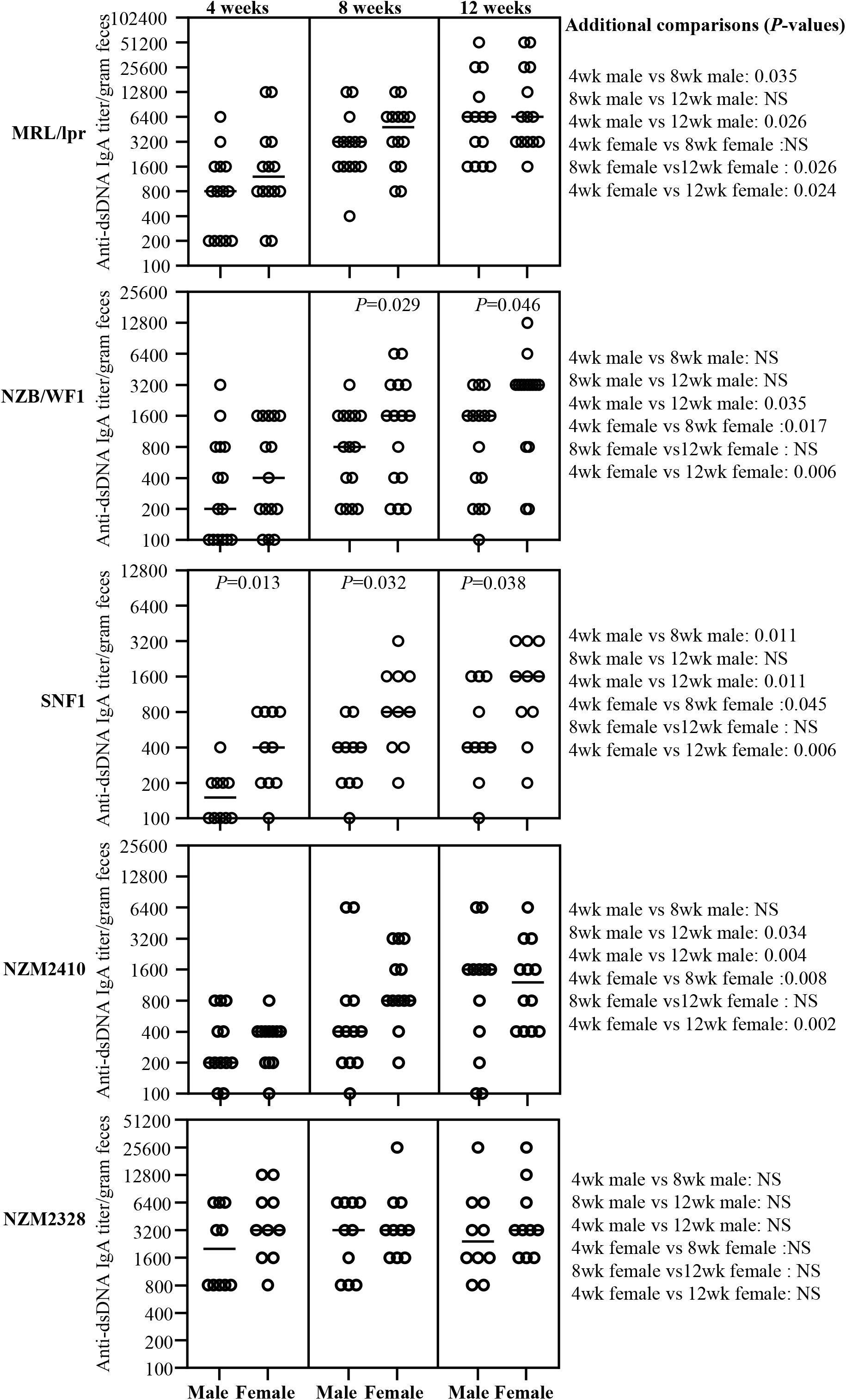
Anti-dsDNA reactive fecal IgA antibodies in lupus-prone mice. Fecal samples were collected from individual mice of indicated strains at 4, 8 and 12 weeks of age and subjected to ELISA to determine dsDNA reactive IgA antibody titer as detailed under materials and method section. dsDNA reactive IgA titers per gram feces are shown. n= 14 males and 14 females for MRL/lpr, 15 males and 15 females for NZB/WF1, 10 males and 10 females for SNF1, 12 males and 12 females for NZM2410, and 10 males and 10 females for NZM2328 strains. *P*-value (by Mann-Whitney test) of male vs female comparisons are shown within the graph. *P*-values of other comparisons are shown on the right.

**Figure 3:**
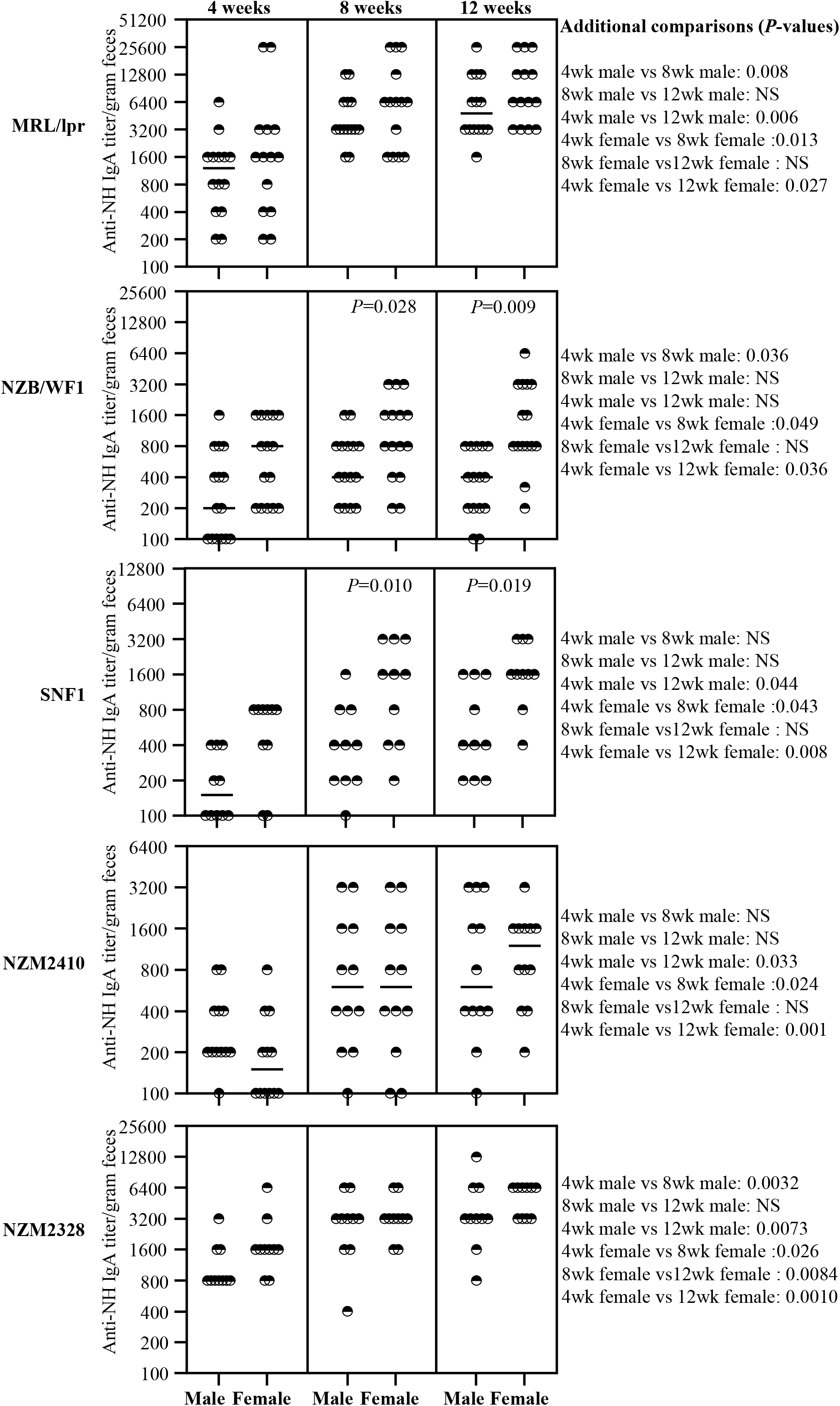
Anti-nucleohistone reactive fecal IgA antibodies in lupus-prone mice. Fecal samples were collected from individual mice of indicated strains at 4, 8 and 12 weeks of age and subjected to ELISA to determine nucleohistone reactive IgA antibody titer as detailed under materials and method section. Nucleohistone reactive IgA titers per gram feces are shown. n= 14 males and 14 females for MRL/lpr, 15 males and 15 females for NZB/WF1, 10 males and 10 females for SNF1, 12 males and 12 females for NZM2410, and 10 males and 10 females for NZM2328 strains. *P*-value (by Mann-Whitney test) of male vs female comparisons are shown within the graph. *P*-values of other comparisons are shown on the right.

### nAg reactivity of fecal IgA of pre-seropositive age correlates with seropositive age circulating autoantibody levels

Previous reports [13, 26, 27, 30, 31] and our results (supplemental Fig. 1) show that significant gender bias in clinical stage kidney disease, as indicated by high proteinuria, is detectable in NZB/WF1, SNF1 and NZM2328 mice, but not in MRL/lpr or NZM2410. On the other hand, gender difference in circulating autoantibody levels is observed primarily in NZB/WF1 and SNF1 mice **(Fig. 4)**. While NZM2328 mice show gender bias in kidney disease, serum autoantibody levels, as reported by others before[27], are comparable in males and females. On the other hand, MRL/lpr mice show some degree of gender bias in the onset of, percentage of mice with, cutaneous lesions as well as survival[32, 33]. However, the differences in proteinuria onset and kidney pathology are not significantly different. Here, we examined if pre-seropositive stage nAg reactivities of fecal IgA from lupus-prone mouse strains correlates with their circulating IgG autoantibody levels at seropositive age. **Fig. 5** shows significant correlations between the pre-seropositive age nAg reactive fecal IgA titers and seropositive age circulating IgG autoantibody levels in both male and female MRL/lpr, NZB/WF1, and SNF1 mice. Although similar trends were observed in NZM2410 and NZM2328 mice, statistically significant correlations were observed only in females and males respectively of these strains. Nevertheless, overall, these results suggest that the pre-seropositive nAg reactivity of fecal IgA titers could be indicative of the degree of eventual systemic autoantibody production in lupus-prone mouse models.

**Figure 4:**
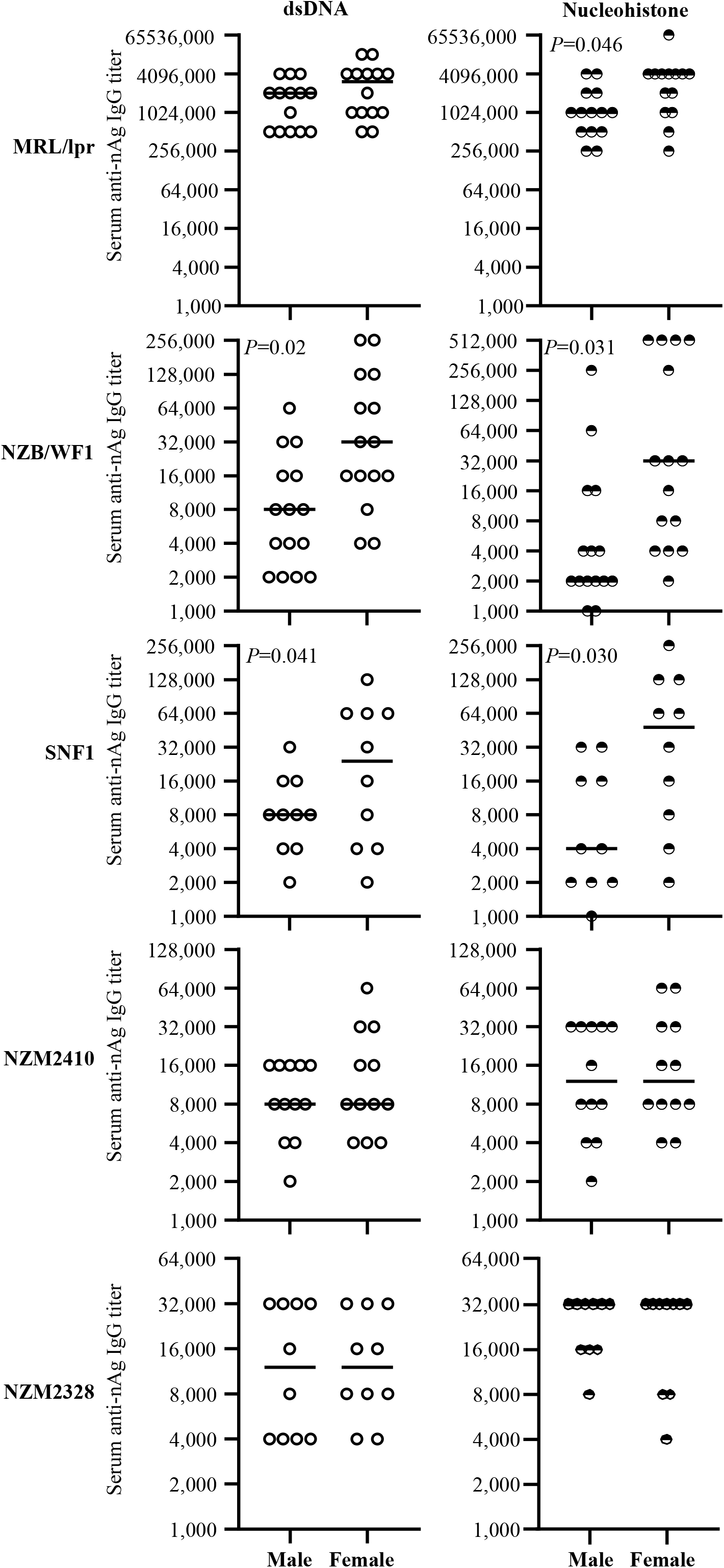
Circulating anti-nAg IgG antibody levels in lupus-prone mouse strains. Serum samples were collected from individual mice of indicated strains at different time-points and subjected to ELISA to determine dsDNA- and nucleohistone- reactive IgG titers as detailed under materials and methods section. nAg-reactive IgG titer of serum samples collected at 12 weeks of age (for MRL/lpr mice) and 16 weeks of age (for all other strains of mice) are shown. n= 14 males and 14 females for MRL/lpr, 15 males and 15 females for NZB/WF1, 10 males and 10 females for SNF1, 12 males and 12 females for NZM2410, and 10 males and 10 females for NZM2328 strains. *P*-value by Mann-Whitney test.

**Figure 5:**
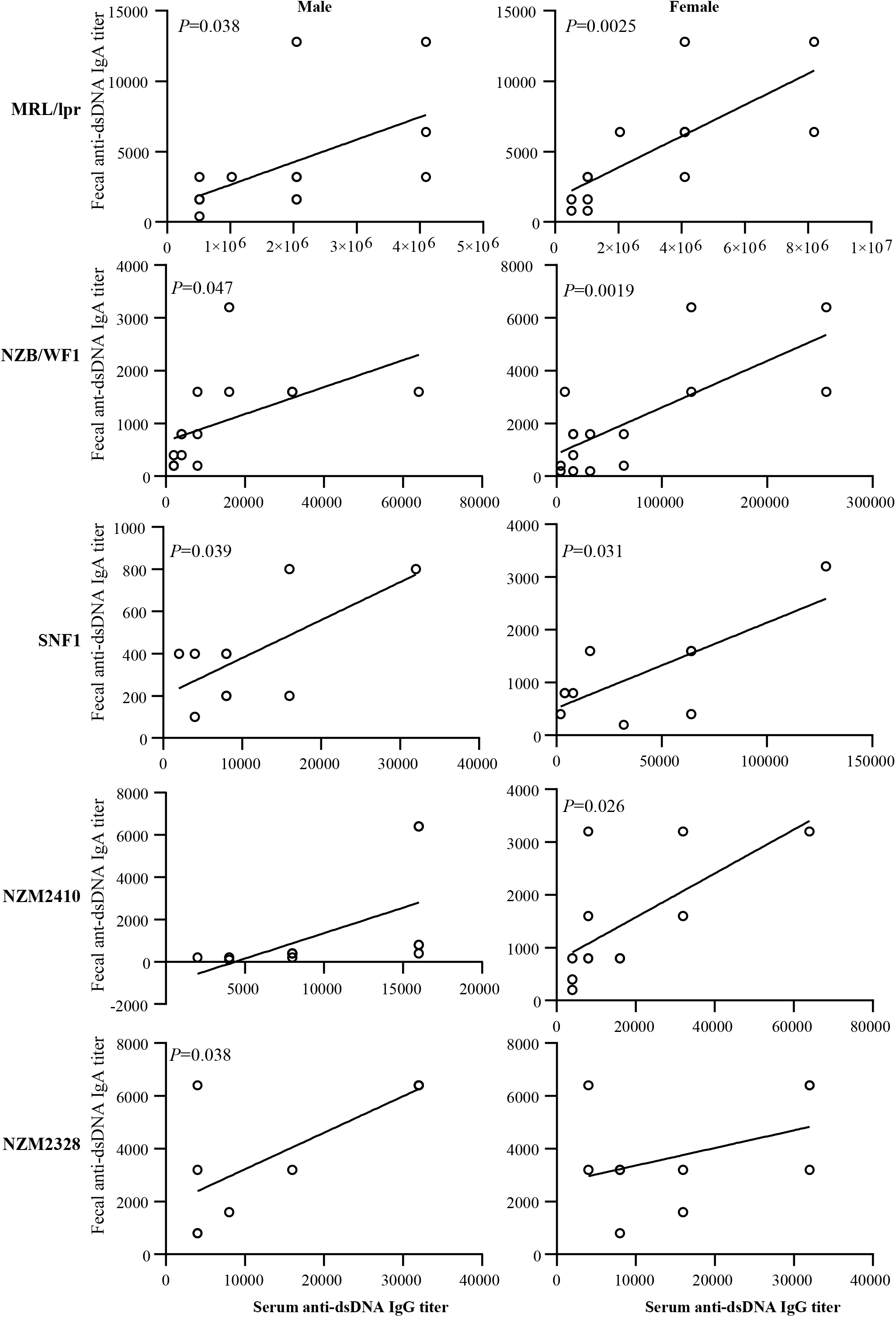
Correlation between pre-seropositive age dsDNA reactivity of fecal IgA and seropositive age circulating anti-dsDNA antibody levels of lupus-prone mice. dsDNA reactive fecal IgA titers were determined using samples collected at 8 weeks of age as described in Fig. 2. Serum anti-dsDNA IgG antibody titers were determined using samples collected at 12 weeks (for MRL/lpr mice) or 16 weeks (for all other indicated strains) as described in supplemental figure 4. Fecal dsDNA reactive IgA titers were correlated with the serum anti-dsDNA IgG titers by Pearson’s correlation approach. n= 14 males and 14 females for MRL/lpr, 15 males and 15 females for NZB/WF1, 10 males and 10 females for SNF1, 12 males and 12 females for NZM2410, and 10 males and 10 females for NZM2328 strains. Similar correlations were detected when nucleohistone reactive IgA titers were compared (not shown).

## Discussion

Previous reports have shown that the degree of IgA production in the gut under normal and clinical conditions including SLE can be different [5, 34, 35]. Fecal IgA levels were found to be higher in SLE patients compared to healthy controls [35]. Using SNF1 and B6 mice, we showed higher amounts of IgA antibody in the fecal samples under lupus-susceptibility [14]. This report also showed that the abundance and nAg reactivities of fecal IgA of SNF1 mice at younger ages correlates positively with rapid disease progression and higher disease incidence of females. Further, we have shown that the immune phenotype including abundance of plasma cells in the gut mucosa is significantly different in male and female SNF1 mice as early as at juvenile age[13, 14, 24]. Here we show that the pre-clinical stage fecal IgA abundance and/or nAg reactivity of lupus-prone mouse strains, MRL/lpr, NZM2328, NZM2410, NZB/WF1 are, similar to SNF1 mice, indicative of the eventual systemic autoantibody levels.

SNF1, NZB/WF1 and NZM2328 mice, all of which show slow disease progression, are known to present strong gender bias in proteinuria onset [13, 26, 27, 30–33]. However, only SNF1 and NZB/WF1, but not NZM2328, showed a difference in the serum levels of nAg reactive antibodies. NZM2410 strain, which was derived from NZB/WF1xNZBF1-crossing, similar to NZM2328, does not present a significant gender bias in disease onset. MRL/lpr mice produce large amounts of nAg reactive antibodies as well as develop skin lesions. This strain also develops lymphoproliferative conditions and kidney disease at much younger ages compared to other strains, and the gender bias in kidney disease is not prominent in this strain. Here, we show that, while the abundance and age-dependent increase in fecal IgA varied among lupus-prone stains of mice, all strains showed considerably higher levels of nAg reactive fecal IgA starting at as early as juvenile age when compared to non-susceptible strain. Further, an overall increase in the abundance and nAg reactivity of fecal IgA can be detected in lupus-prone mice at adult ages as compared to juvenile age. We also show that nAg reactivity of fecal IgA at younger age correlates with the eventual systemic autoantibody levels as well as known gender bias in disease incidence in most lupus-prone strains.

As evident from the literature, gender bias in the autoantibody production and clinical disease onset are not ubiquitous among lupus-prone mouse strains. SNF1 and NZB/WF1 strains show clear differences in autoantibody production and the timing of clinical disease onset among males and females[13, 31]. Our results show that statistically significant differences in the fecal IgA abundance and nAg reactivity between males and females are found only in these two out of five strains of mice tested. Interestingly, although strong gender bias is observed in the kidney disease incidence in NZM2328 mice, systemic autoantibody levels are comparable in males and females of this strain[27]. We found that, similar to serum autoantibody levels, fecal IgA abundance and nAg reactivity are not profoundly different among male and female NZM2328 mice. Overall, our observations indicate that the magnitude of nAg reactive antibody production in the gut and systemic compartment are similar in lupus prone backgrounds but nAg reactive IgA antibodies are detected in the feces relatively earlier.

Pro-inflammatory features including hyper-activation of B cells of the gut mucosa, in the presence of gut microbial components, could aid in the production of anti-microbial antibodies that cross-react with host nAg. In fact, although it is not known if all lupus-prone mice show similar features, our studies using SNF1 mice, which presents a strong gender bias in disease, showed that female gut mucosa is more inflamed than that of males at as early as juvenile age[13, 24, 36]. Importantly, gut microbiota including symbionts and pathobionts can influence the magnitude of antibody production in the gut mucosa and the IgA abundance in feces [37–41]. However, previous reports, including ours, have shown that gut microbiota composition is significantly different in autoimmune prone males and females only at adult age [24, 42, 43]. Therefore, it is possible that a combination of gut proinflammatory events, but not the microbiota composition itself, is responsible for differences in the fecal IgA features in males and females of some strains of mice. Irrespective of the differences in gut microbiota composition, microbial antigens are continuously sampled by immune cells of the gut mucosa and it can result in local and systemic responses [44–47]. Importantly, microbial nucleic acids and histone-like proteins can be structurally homologous to their mammalian host counterparts [48–52]. Therefore, gender specific gut immune features of each strain, in combination with microbial factors that induce antibody production, could determine its fecal IgA features including the degree of nAg reactivity. Our observation that fecal IgA in lupus-prone mice is dsDNA and nucleohistone reactive compared to lupus-resistant mice at as early as juvenile age also suggests that gut mucosa could be the initial site of autoantibody production under lupus susceptibility. An age-dependent increase in the dsDNA and nucleohistone reactive fecal IgA levels in lupus-prone mouse strains also supports this notion.

Overall, our study demonstrates that fecal IgA profiles vary in different strains of lupus-prone mice, but their nAg-reactivity shows a trend similar to that of the circulating autoantibody levels. Importantly, nAg reactivity of fecal IgA is detectable at as early as juvenile age in all lupus-prone mouse strains. Furthermore, the younger age fecal IgA features are, in most part, reflective of eventual circulating autoantibody levels and the known gender bias in disease onset. These observations, in association with our recent report[14], suggest that fecal IgA features, nuclear antigen reactivity particularly, could be employed to detect the systemic autoimmune activities in at-risk subjects, before the seropositivity and clinical stage disease onset. Hence, further studies employing longitudinal samples from SLE at-risk subjects are needed to validate this notion and to confirm the biomarker value of fecal IgA features in predicting systemic autoimmunity.

## *Acknowledgments

This work was supported by internal funds from MUSC, National Institutes of Health (NIH) grants R21AI136339 and R01AI138511. R.G. performed experiments and reviewed the paper, S.R. performed experiments and reviewed the manuscript, W.S. performed experiments, and C.V. designed the study, performed the experiments, and wrote the paper. Dr. Vasu is the guarantor of this work and, as such, had full access to all the data in the study and takes responsibility for the integrity and accuracy of the data and analysis.

## References

[1] R. Gualtierotti, M. Biggioggero, A. E. Penatti, P. L. Meroni. Updating on the pathogenesis of systemic lupus erythematosus. Autoimmun Rev, 2010 10: 3–7.

[2] L. Li, C. Mohan. Genetic basis of murine lupus nephritis. Semin Nephrol, 2007;27: 12–21.

[3] D. Webber, J. Cao, D. Dominguez, D. D. Gladman, D. M. Levy, L. Ng et al. Association of systemic lupus erythematosus (SLE) genetic susceptibility loci with lupus nephritis in childhood-onset and adult-onset SLE. Rheumatology, 2019.

[4] S. K. Tedeschi, B. Bermas, K. H. Costenbader. Sexual disparities in the incidence and course of SLE and RA. Clin Immunol, 2013;149: 211–8.

[5] D. Azzouz, A. Omarbekova, A. Heguy, D. Schwudke, N. Gisch, B. H. Rovin et al. Lupus nephritis is linked to disease-activity associated expansions and immunity to a gut commensal. Ann Rheum Dis, 2019;78: 947–56.

[6] D. F. Zegarra-Ruiz, A. El Beidaq, A. J. Iniguez, M. Lubrano Di Ricco, S. Manfredo Vieira, W. E. Ruff et al. A Diet-Sensitive Commensal Lactobacillus Strain Mediates TLR7-Dependent Systemic Autoimmunity. Cell Host Microbe, 2019;25: 113–27 e6.

[7] X. M. Luo, M. R. Edwards, Q. Mu, Y. Yu, M. D. Vieson, C. M. Reilly et al. Gut Microbiota in Human Systemic Lupus Erythematosus and a Mouse Model of Lupus. Applied and environmental microbiology, 2018;84.

[8] P. Lopez, B. de Paz, J. Rodriguez-Carrio, A. Hevia, B. Sanchez, A. Margolles et al. Th17 responses and natural IgM antibodies are related to gut microbiota composition in systemic lupus erythematosus patients. Sci Rep, 2016;6: 24072.

[9] B. M. Johnson, M. C. Gaudreau, M. M. Al-Gadban, R. Gudi, C. Vasu. Impact of dietary deviation on disease progression and gut microbiome composition in lupus-prone SNF1 mice. Clin Exp Immunol, 2015;181: 323–37.

[10] P. J. Turnbaugh, V. K. Ridaura, J. J. Faith, F. E. Rey, R. Knight, J. I. Gordon. The effect of diet on the human gut microbiome: a metagenomic analysis in humanized gnotobiotic mice. Sci Transl Med, 2009;1: 6ra14.

[11] M. R. Edwards, R. Dai, B. Heid, T. E. Cecere, D. Khan, Q. Mu et al. Commercial rodent diets differentially regulate autoimmune glomerulonephritis, epigenetics and microbiota in MRL/lpr mice. Int Immunol, 2017;29: 263–76.

[12] B. M. Johnson, M. C. Gaudreau, R. Gudi, R. Brown, G. Gilkeson, C. Vasu. Gut microbiota differently contributes to intestinal immune phenotype and systemic autoimmune progression in female and male lupus-prone mice. J Autoimmun, 2020;108: 102420.

[13] M. C. Gaudreau, B. M. Johnson, R. Gudi, M. M. Al-Gadban, C. Vasu. Gender bias in lupus: does immune response initiated in the gut mucosa have a role? Clinical and experimental immunology, 2015;180: 393–407.

[14] W. Sun, R. R. Gudi, B. M. Johnson, C. Vasu. Abundance and nuclear antigen reactivity of intestinal and fecal Immunoglobulin A in lupus-prone mice at younger ages correlate with the onset of eventual systemic autoimmunity. Sci Rep, 2020;10: 14258.

[15] A. Nakajima, A. Vogelzang, M. Maruya, M. Miyajima, M. Murata, A. Son et al. IgA regulates the composition and metabolic function of gut microbiota by promoting symbiosis between bacteria. The Journal of experimental medicine, 2018;215: 2019–34.

[16] J. Fadlallah, H. El Kafsi, D. Sterlin, C. Juste, C. Parizot, K. Dorgham et al. Microbial ecology perturbation in human IgA deficiency. Sci Transl Med, 2018;10.

[17] G. P. Donaldson, M. S. Ladinsky, K. B. Yu, J. G. Sanders, B. B. Yoo, W. C. Chou et al. Gut microbiota utilize immunoglobulin A for mucosal colonization. Science, 2018;360: 795–800.

[18] D. Villalta, N. Bizzaro, N. Bassi, M. Zen, M. Gatto, A. Ghirardello et al. Anti-dsDNA antibody isotypes in systemic lupus erythematosus: IgA in addition to IgG anti-dsDNA help to identify glomerulonephritis and active disease. PloS one, 2013;8: e71458.

[19] Y. Jia, L. Zhao, C. Wang, J. Shang, Y. Miao, Y. Dong et al. Anti-Double-Stranded DNA Isotypes and Anti-C1q Antibody Improve the Diagnostic Specificity of Systemic Lupus Erythematosus. Dis Markers, 2018;2018: 4528547.

[20] M. Gripenberg, T. Helve. Anti-DNA antibodies of IgA class in patients with systemic lupus erythematosus. Rheumatology international, 1986;6: 53–5.

[21] S. A. Jost, L. C. Tseng, L. A. Matthews, R. Vasquez, S. Zhang, K. B. Yancey et al. IgG, IgM, and IgA antinuclear antibodies in discoid and systemic lupus erythematosus patients. ScientificWorldJournal, 2014;2014: 171028.

[22] A. M. Miltenburg, A. Roos, L. Slegtenhorst, M. R. Daha, F. C. Breedveld. IgA anti-dsDNA antibodies in systemic lupus erythematosus: occurrence, incidence and association with clinical and laboratory variables of disease activity. The Journal of rheumatology, 1993;20: 53–8.

[23] N. R. Gompertz, D. A. Isenberg, B. M. Turner. Correlation between clinical features of systemic lupus erythematosus and levels of antihistone antibodies of the IgG, IgA, and IgM isotypes. Ann Rheum Dis, 1990;49: 524–7.

[24] G. M. Johnson BM, Gudi R, Brown R, Gilkeson G, and Vasu C. Gut microbiota differently contributes to intestinal immune phenotype and systemic autoimmune progression in female and male lupus-prone mice. BioRxiv 2019.

[25] U. H. Rudofsky, D. A. Lawrence. New Zealand mixed mice: a genetic systemic lupus erythematosus model for assessing environmental effects. Environ Health Perspect, 1999;107 Suppl 5: 713–21.

[26] U. H. Rudofsky, B. D. Evans, S. L. Balaban, V. D. Mottironi, A. E. Gabrielsen. Differences in expression of lupus nephritis in New Zealand mixed H-2z homozygous inbred strains of mice derived from New Zealand black and New Zealand white mice. Origins and initial characterization. Laboratory investigation; a journal of technical methods and pathology, 1993;68: 419–26.

[27] S. T. Waters, S. M. Fu, F. Gaskin, U. S. Deshmukh, S. S. Sung, C. C. Kannapell et al. NZM2328: a new mouse model of systemic lupus erythematosus with unique genetic susceptibility loci. Clin Immunol, 2001;100: 372–83.

[28] L. Morel. Mapping lupus susceptibility genes in the NZM2410 mouse model. Adv Immunol, 2012;115: 113–39.

[29] T. Celhar, A. M. Fairhurst. Modelling clinical systemic lupus erythematosus: similarities, differences and success stories. Rheumatology, 2017;56: i88–i99.

[30] M. L. Richard, G. Gilkeson. Mouse models of lupus: what they tell us and what they don’t. Lupus Sci Med, 2018;5: e000199.

[31] B. S. Andrews, R. A. Eisenberg, A. N. Theofilopoulos, S. Izui, C. B. Wilson, P. J. McConahey et al. Spontaneous murine lupus-like syndromes. Clinical and immunopathological manifestations in several strains. The Journal of experimental medicine, 1978;148: 1198–215.

[32] J. L. Goulet, R. C. Griffiths, P. Ruiz, R. F. Spurney, D. S. Pisetsky, B. H. Koller et al. Deficiency of 5- lipoxygenase abolishes sex-related survival differences in MRL-lpr/lpr mice. Journal of immunology, 1999;163: 359–66.

[33] H. Kanauchi, F. Furukawa, S. Imamura. Characterization of cutaneous infiltrates in MRL/lpr mice monitored from onset to the full development of lupus erythematosus-like skin lesions. J Invest Dermatol, 1991;96: 478–83.

[34] M. Dzidic, T. R. Abrahamsson, A. Artacho, B. Bjorksten, M. C. Collado, A. Mira et al. Aberrant IgA responses to the gut microbiota during infancy precede asthma and allergy development. J Allergy Clin Immunol, 2017;139: 1017–25 e14.

[35] L. Frehn, A. Jansen, E. Bennek, A. D. Mandic, I. Temizel, S. Tischendorf et al. Distinct patterns of IgG and IgA against food and microbial antigens in serum and feces of patients with inflammatory bowel diseases. PloS one, 2014;9: e106750.

[36] S. Esposito, S. Bosis, M. Semino, D. Rigante. Infections and systemic lupus erythematosus. Eur J Clin Microbiol Infect Dis, 2014;33: 1467–75.

[37] J. D. Planer, Y. Peng, A. L. Kau, L. V. Blanton, I. M. Ndao, P. I. Tarr et al. Development of the gut microbiota and mucosal IgA responses in twins and gnotobiotic mice. Nature, 2016;534: 263–6.

[38] D. A. Peterson, N. P. McNulty, J. L. Guruge, J. I. Gordon. IgA response to symbiotic bacteria as a mediator of gut homeostasis. Cell Host Microbe, 2007;2: 328–39.

[39] S. Hapfelmeier, M. A. Lawson, E. Slack, J. K. Kirundi, M. Stoel, M. Heikenwalder et al. Reversible microbial colonization of germ-free mice reveals the dynamics of IgA immune responses. Science, 2010;328: 1705–9.

[40] E. Lecuyer, S. Rakotobe, H. Lengline-Garnier, C. Lebreton, M. Picard, C. Juste et al. Segmented filamentous bacterium uses secondary and tertiary lymphoid tissues to induce gut IgA and specific T helper 17 cell responses. Immunity, 2014;40: 608–20.

[41] A. J. Macpherson, L. Hunziker, K. McCoy, A. Lamarre. IgA responses in the intestinal mucosa against pathogenic and non-pathogenic microorganisms. Microbes Infect, 2001;3: 1021–35.

[42] L. Yurkovetskiy, M. Burrows, A. A. Khan, L. Graham, P. Volchkov, L. Becker et al. Gender bias in autoimmunity is influenced by microbiota. Immunity, 2013;39: 400–12.

[43] J. G. Markle, D. N. Frank, S. Mortin-Toth, C. E. Robertson, L. M. Feazel, U. Rolle-Kampczyk et al. Sex differences in the gut microbiome drive hormone-dependent regulation of autoimmunity. Science, 2013;339: 1084–8.

[44] D. Rios, M. B. Wood, J. Li, B. Chassaing, A. T. Gewirtz, I. R. Williams. Antigen sampling by intestinal M cells is the principal pathway initiating mucosal IgA production to commensal enteric bacteria. Mucosal Immunol, 2016;9: 907–16.

[45] O. Schulz, O. Pabst. Antigen sampling in the small intestine. Trends in immunology, 2013;34: 155–61.

[46] A. J. Stagg, A. L. Hart, S. C. Knight, M. A. Kamm. The dendritic cell: its role in intestinal inflammation and relationship with gut bacteria. Gut, 2003;52: 1522–9.

[47] Q. Zhao, C. O. Elson. Adaptive immune education by gut microbiota antigens. Immunology, 2018;154: 28–37.

[48] G. S. Gilkeson, J. P. Grudier, D. G. Karounos, D. S. Pisetsky. Induction of anti-double stranded DNA antibodies in normal mice by immunization with bacterial DNA. Journal of immunology, 1989;142: 1482–6.

[49] D. S. Pisetsky, J. P. Grudier, G. S. Gilkeson. A role for immunogenic DNA in the pathogenesis of systemic lupus erythematosus. Arthritis and rheumatism, 1990;33: 153–9.

[50] A. Sabbatini, S. Bombardieri, P. Migliorini. Autoantibodies from patients with systemic lupus erythematosus bind a shared sequence of SmD and Epstein-Barr virus-encoded nuclear antigen EBNA I. European journal of immunology, 1993;23: 1146–52.

[51] A. Balandina, D. Kamashev, J. Rouviere-Yaniv. The bacterial histone-like protein HU specifically recognizes similar structures in all nucleic acids. DNA, RNA, and their hybrids. The Journal of biological chemistry, 2002;277: 27622–8.

[52] D. Kamashev, Y. Agapova, S. Rastorguev, A. A. Talyzina, K. M. Boyko, D. A. Korzhenevskiy et al. Comparison of histone-like HU protein DNA-binding properties and HU/IHF protein sequence alignment. PloS one, 2017;12: e0188037.

