## supplemental figures for "Preclinical stage abundance and nuclear antigen reactivity of fecal Immunoglobulin A (IgA) varies among males and females of lupus-prone mouse models"

Fig. 1

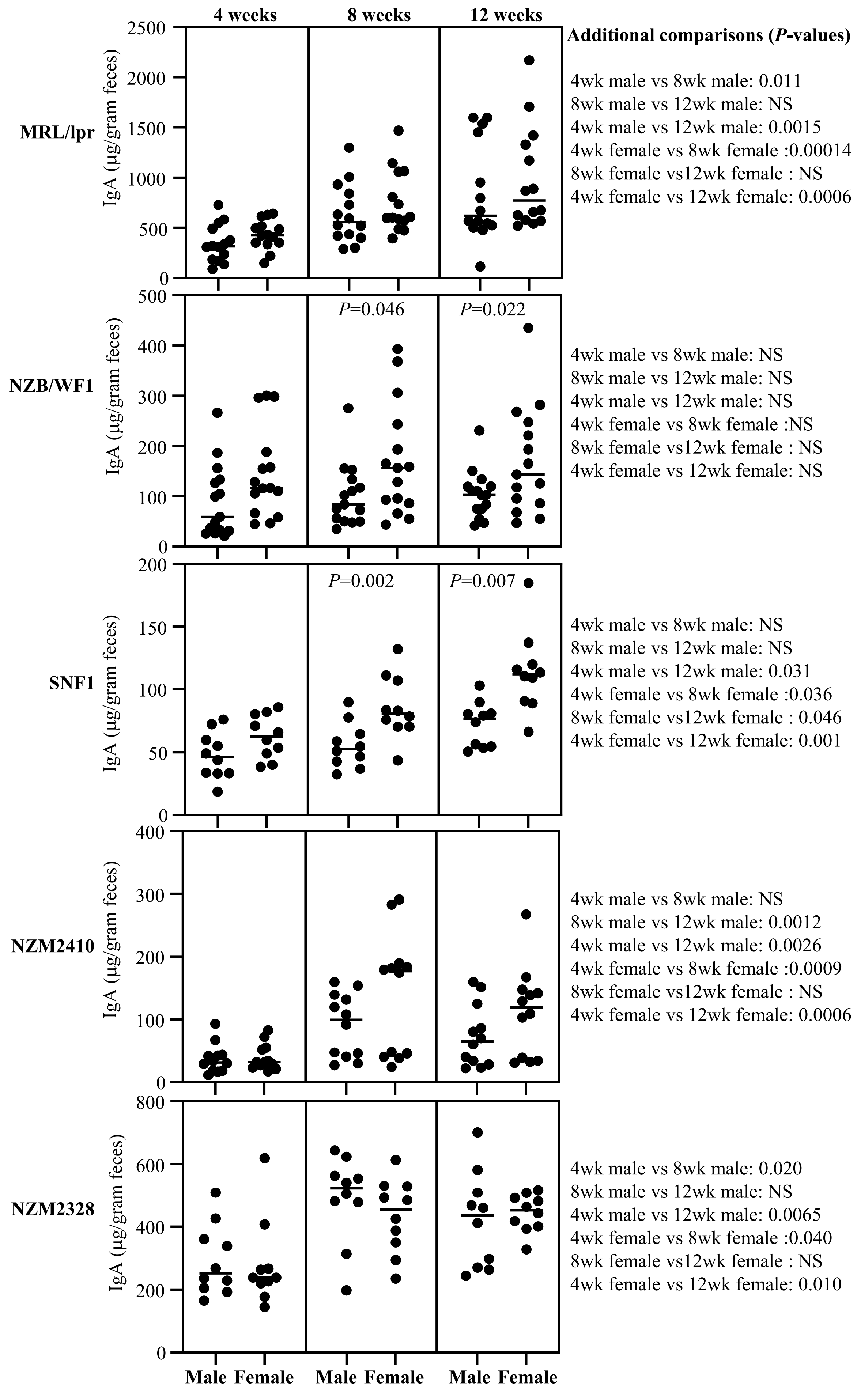
4 weeks
8 weeks
12 weeks
Additional comparisons (*P*-values)
MRL/lpr
IgA (µg/gram feces)
2500
2000
1500
1000
500
0
4wk male vs 8wk male: 0.011
8wk male vs 12wk male: NS
4wk male vs 12wk male: 0.0015
4wk female vs 8wk female :0.00014
8wk female vs12wk female : NS
4wk female vs 12wk female: 0.0006
NZB/WF1
IgA (µg/gram feces)
500
400
300
200
100
0
*P*=0.046
*P*=0.022
4wk male vs 8wk male: NS
8wk male vs 12wk male: NS
4wk male vs 12wk male: NS
4wk female vs 8wk female :NS
8wk female vs12wk female : NS
4wk female vs 12wk female: NS
SNF1
IgA (µg/gram feces)
200
150
100
50
0
*P*=0.002
*P*=0.007
4wk male vs 8wk male: NS
8wk male vs 12wk male: NS
4wk male vs 12wk male: 0.031
4wk female vs 8wk female :0.036
8wk female vs12wk female : 0.046
4wk female vs 12wk female: 0.001
NZM2410
IgA (µg/gram feces)
400
300
200
100
0
4wk male vs 8wk male: NS
8wk male vs 12wk male: 0.0012
4wk male vs 12wk male: 0.0026
4wk female vs 8wk female :0.0009
8wk female vs12wk female : NS
4wk female vs 12wk female: 0.0006
NZM2328
IgA (µg/gram feces)
800
600
400
200
0
4wk male vs 8wk male: 0.020
8wk male vs 12wk male: NS
4wk male vs 12wk male: 0.0065
4wk female vs 8wk female :0.040
8wk female vs12wk female : NS
4wk female vs 12wk female: 0.010
Male Female
Male Female
Male Female

Fig. 2

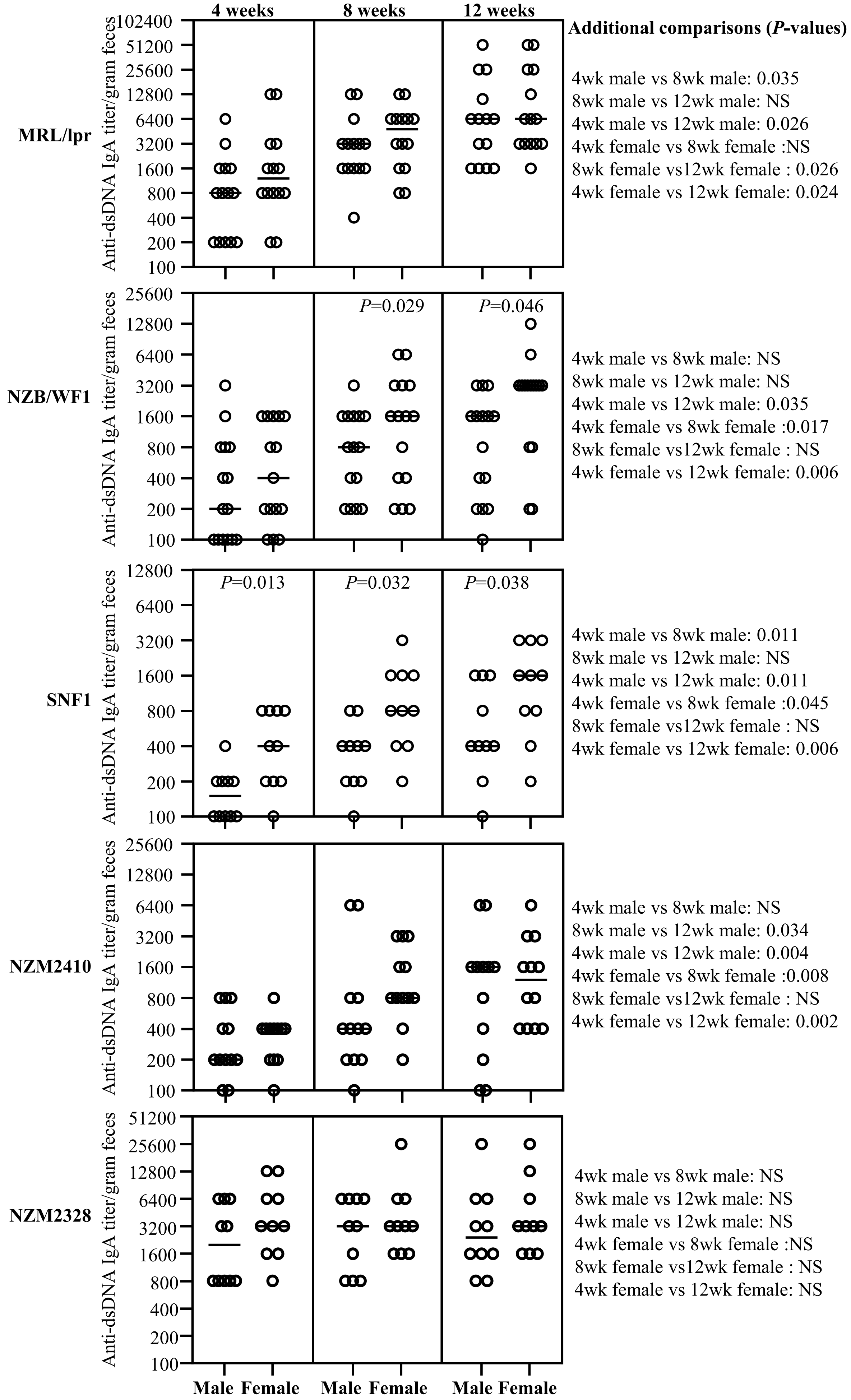
4 weeks
8 weeks
12 weeks
Additional comparisons (P-values)
MRL/lpr
4wk male vs 8wk male: 0.035
8wk male vs 12wk male: NS
4wk male vs 12wk male: 0.026
4wk female vs 8wk female :NS
8wk female vs12wk female : 0.026
4wk female vs 12wk female: 0.024
NZB/WF1
P=0.029
P=0.046
4wk male vs 8wk male: NS
8wk male vs 12wk male: NS
4wk male vs 12wk male: 0.035
4wk female vs 8wk female :0.017
8wk female vs12wk female : NS
4wk female vs 12wk female: 0.006
SNF1
P=0.013
P=0.032
P=0.038
4wk male vs 8wk male: 0.011
8wk male vs 12wk male: NS
4wk male vs 12wk male: 0.011
4wk female vs 8wk female :0.045
8wk female vs12wk female : NS
4wk female vs 12wk female: 0.006
NZM2410
4wk male vs 8wk male: NS
8wk male vs 12wk male: 0.034
4wk male vs 12wk male: 0.004
4wk female vs 8wk female :0.008
8wk female vs12wk female : NS
4wk female vs 12wk female: 0.002
NZM2328
4wk male vs 8wk male: NS
8wk male vs 12wk male: NS
4wk male vs 12wk male: NS
4wk female vs 8wk female :NS
8wk female vs12wk female : NS
4wk female vs 12wk female: NS
Anti-dsDNA IgA titer/gram feces
Male Female
Male Female
Male Female

Fig. 3

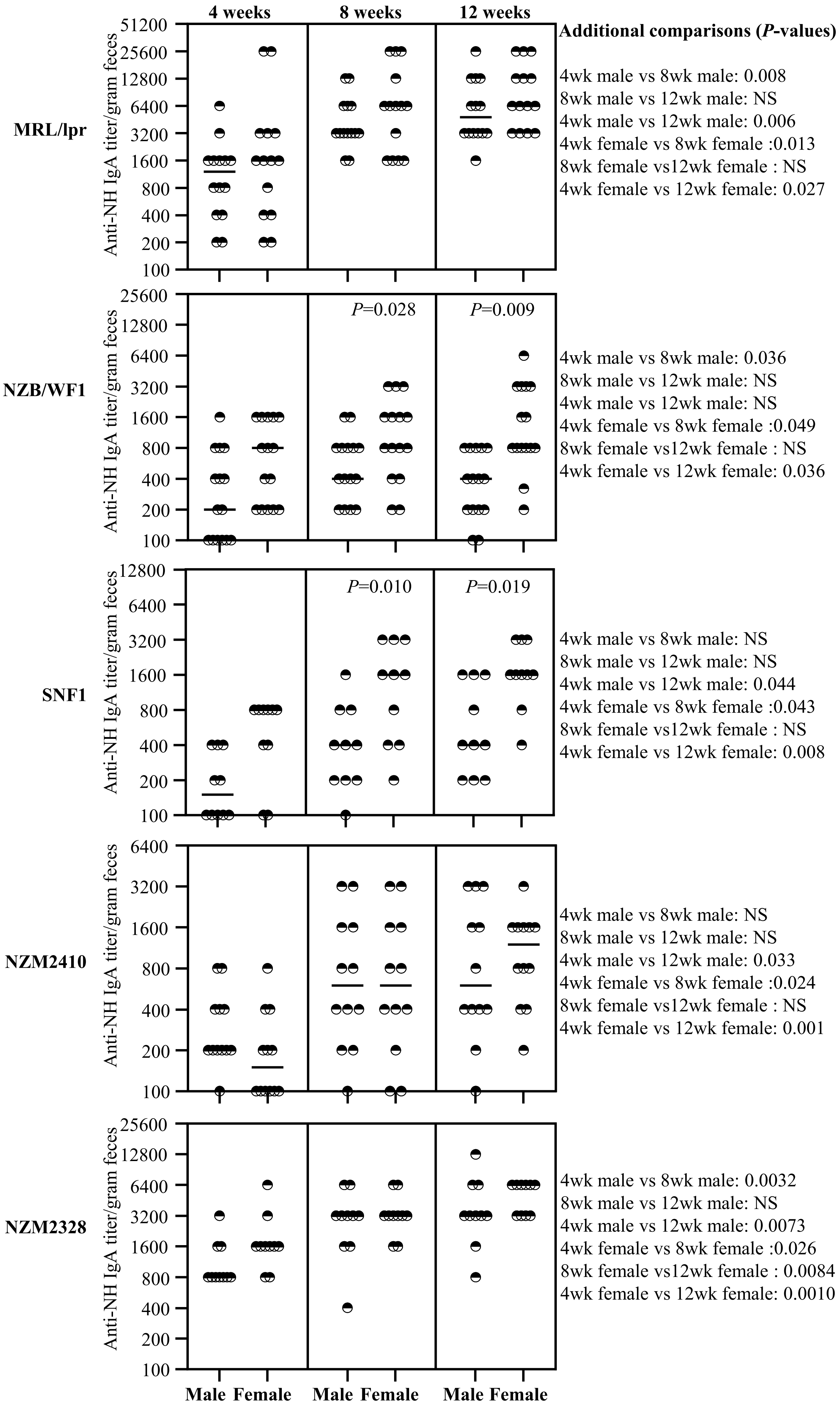
4 weeks
8 weeks
12 weeks
Additional comparisons (P-values)
MRL/lpr
4wk male vs 8wk male: 0.008
8wk male vs 12wk male: NS
4wk male vs 12wk male: 0.006
4wk female vs 8wk female :0.013
8wk female vs12wk female : NS
4wk female vs 12wk female: 0.027
NZB/WF1
P=0.028
P=0.009
4wk male vs 8wk male: 0.036
8wk male vs 12wk male: NS
4wk male vs 12wk male: NS
4wk female vs 8wk female :0.049
8wk female vs12wk female : NS
4wk female vs 12wk female: 0.036
SNF1
P=0.010
P=0.019
4wk male vs 8wk male: NS
8wk male vs 12wk male: NS
4wk male vs 12wk male: 0.044
4wk female vs 8wk female :0.043
8wk female vs12wk female : NS
4wk female vs 12wk female: 0.008
NZM2410
4wk male vs 8wk male: NS
8wk male vs 12wk male: NS
4wk male vs 12wk male: 0.033
4wk female vs 8wk female :0.024
8wk female vs12wk female : NS
4wk female vs 12wk female: 0.001
NZM2328
4wk male vs 8wk male: 0.0032
8wk male vs 12wk male: NS
4wk male vs 12wk male: 0.0073
4wk female vs 8wk female :0.026
8wk female vs12wk female : 0.0084
4wk female vs 12wk female: 0.0010
Anti-NH IgA titer/gram feces
51200
25600
12800
6400
3200
1600
800
400
200
100
Male Female Male Female Male Female

Fig. 4

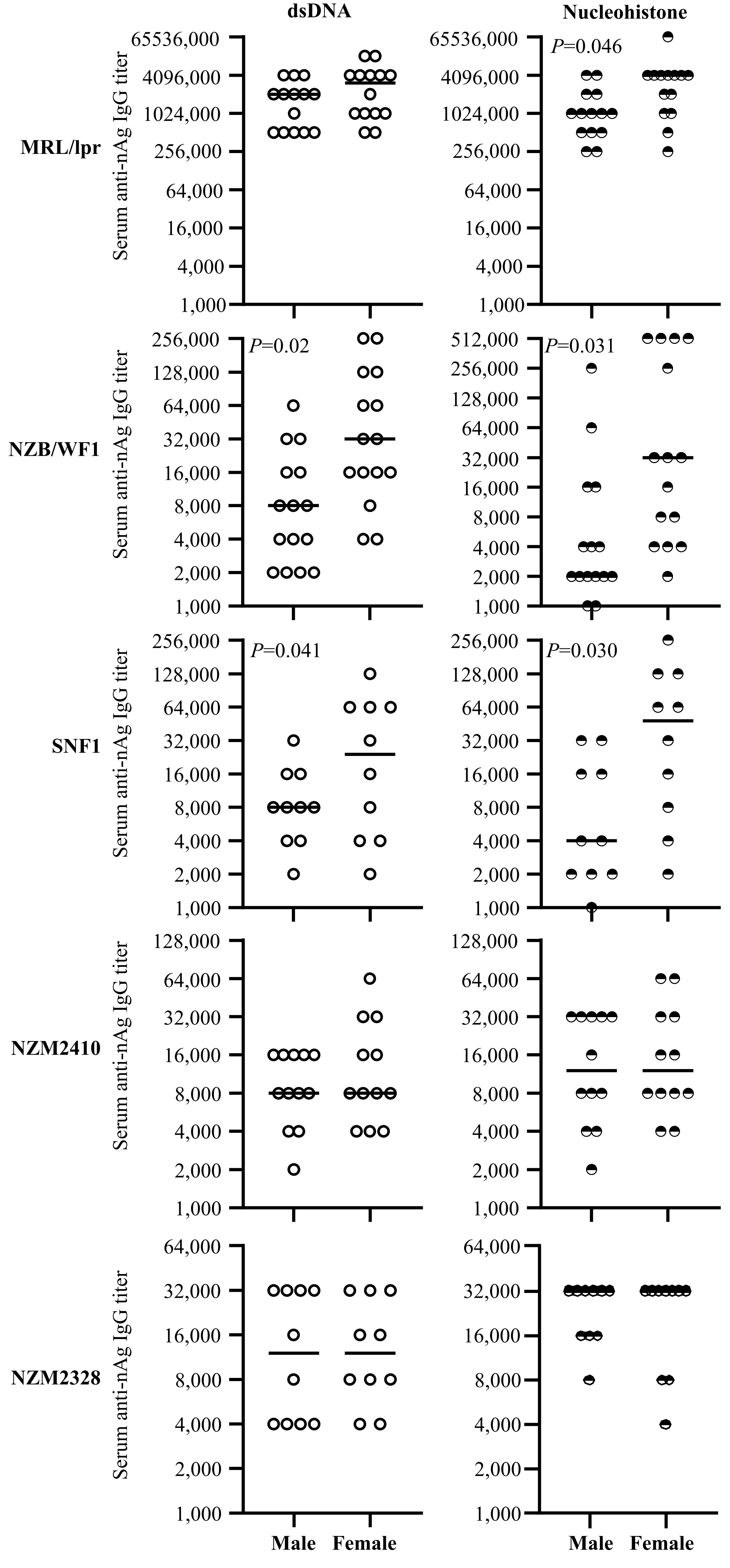
dsDNA
Nucleohistone
MRL/lpr
NZB/WF1
SNF1
NZM2410
NZM2328
Serum anti-nAg IgG titer
P=0.046
P=0.02
P=0.031
P=0.041
P=0.030
65536,000
4096,000
1024,000
512,000
256,000
128,000
64,000
32,000
16,000
8,000
4,000
2,000
1,000
Male
Female

Fig. 5

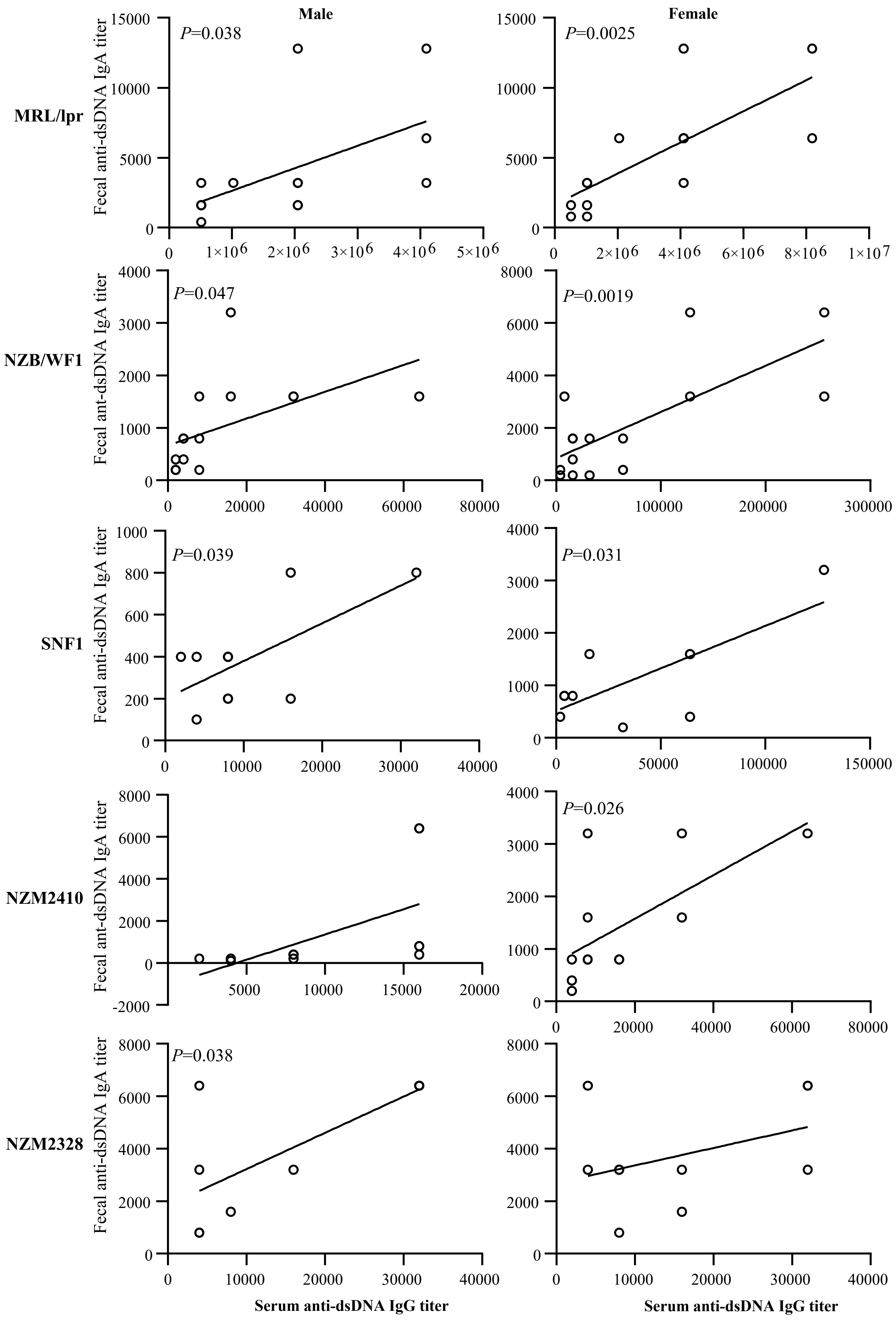
Male
Female
MRL/lpr
NZB/WF1
SNF1
NZM2410
NZM2328
P=0.038
P=0.0025
P=0.047
P=0.0019
P=0.039
P=0.031
P=0.026
P=0.038
Fecal anti-dsDNA IgA titer
Fecal ant-dsDNA IgA titer
Serum anti-dsDNA IgG titer
Serum anti-dsDNA IgG titer

Supplemental Fig. 1

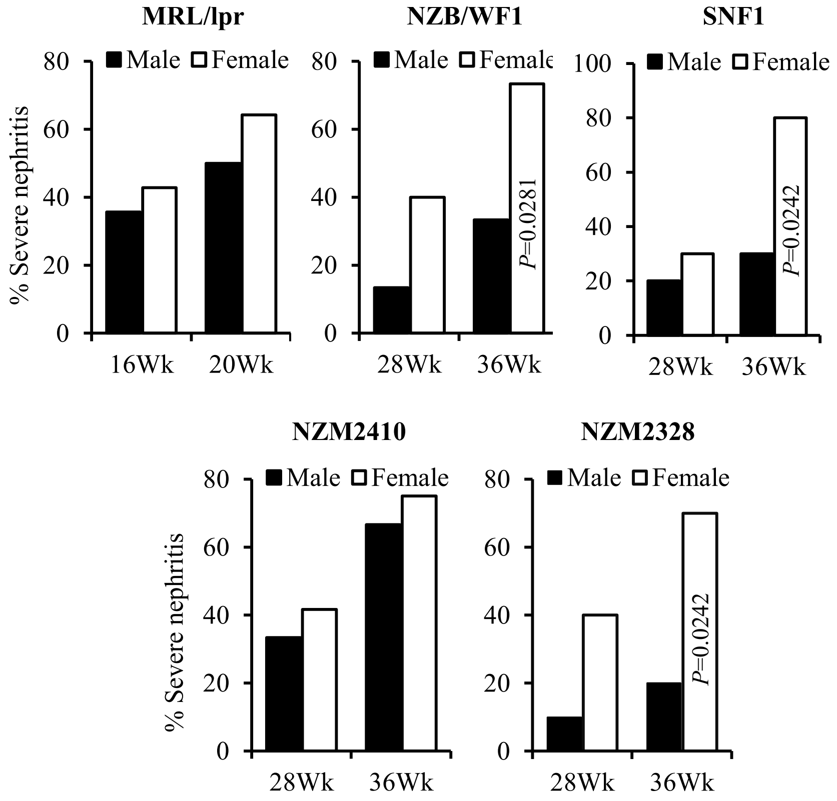


**Supplemental figure 1: Differences in severe nephritis incidence in lupus-prone male and female mice.** Urine samples were collected from all mice used for this study at timely intervals and tested for protein levels as described under the materials and methods section. Mice that showed proteinuria (>5 mg/ml) were considered to have severe nephritis. n= 14 males and 14 females for MRL/lpr, 15 males and 15 females for NZB/WF1, 10 males and 10 females for SNF1, 12 males and 12 females for NZM2410, and 10 males and 10 females for NZM2328 strains. *P*-value (by Mann-Whitney test) of specific age male vs female comparisons are shown.

Supplemental Fig. 2

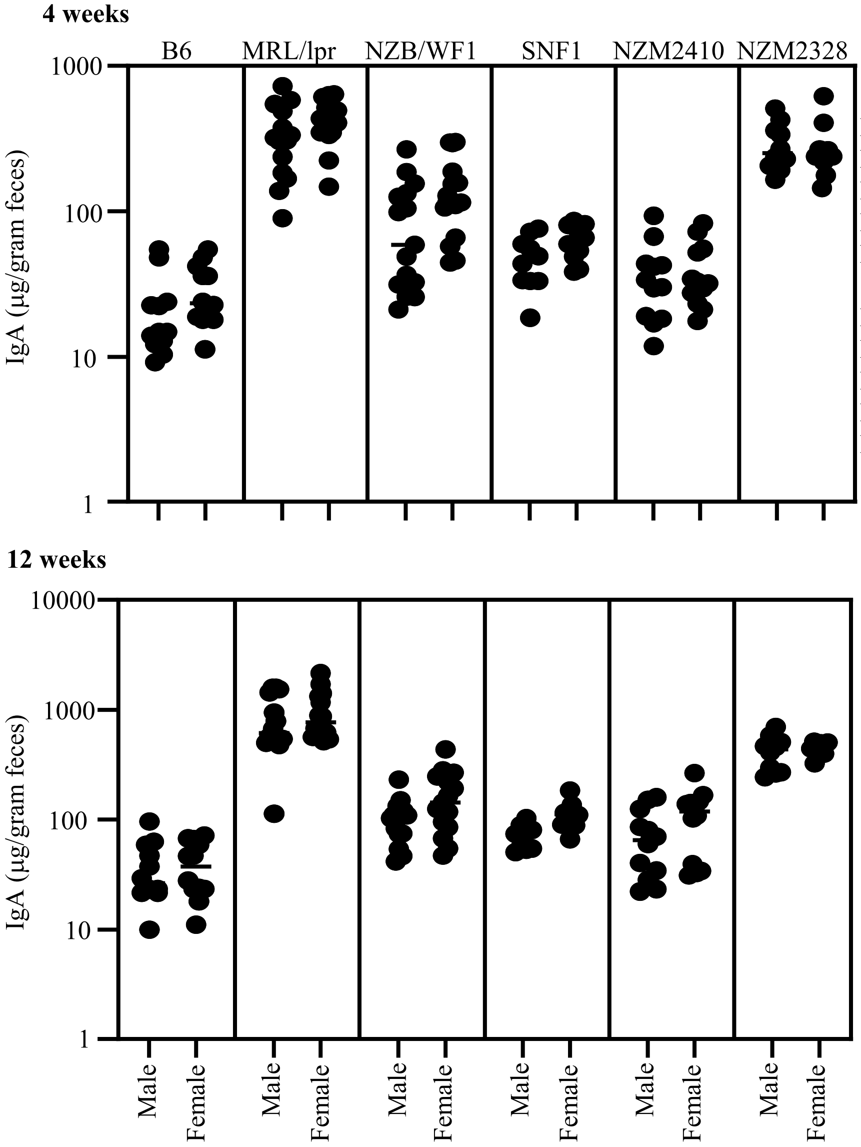


**Comparisons (*P*-values)**

B6 male vs MRL/lpr male: <0.0001
B6 male vs NZB/WF1 male:0.0035
B6 male vs SNF1 male:0.0016
B6 male vs NZM2410 male:NS
B6 male vs NZM2328 male: <0.0001

B6 female vs MRL/lpr female: <0.0001
B6 female vs NZB/WF1 female: 0.00014
B6 female vs SNF1 female: <0.0001
B6 female vs NZM2410 female: NS
B6 female vs NZM2328 female: <0.0001

B6 male vs MRL/lpr male: <0.0001
B6 male vs NZB/WF1 male:0.0002
B6 male vs SNF1 male:0.0014
B6 male vs NZM2410 male:0.034
B6 male vs NZM2328 male: <0.0001

B6 female vs MRL/lpr female: <0.0001
B6 female vs NZB/WF1 female: 0.0003
B6 female vs SNF1 female: <0.0001
B6 female vs NZM2410 female: 0.0029
B6 female vs NZM2328 female: <0.0001

**Supplemental figure 2: Abundance of IgA in the feces of lupus-susceptible and non-susceptible mouse strains.** Fecal samples were collected from individual mice of indicated strains and subjected to ELISA to determine IgA concentrations as detailed under materials and methods section. IgA concentrations per gram feces (collected at 4 and 12 weeks of age) are shown. n= 12 males and 12 females for B6 mice, 4 males and 14 females for MRL/lpr, 15 males and 15 females for NZB/WF1, 10 males and 10 females for SNF1, 12 males and 12 females for NZM2410, and 10 males and 10 females for NZM2328 strains. *P*-value (by Mann-Whitney test) of various comparisons are shown on the right.

Supplemental Fig. 3

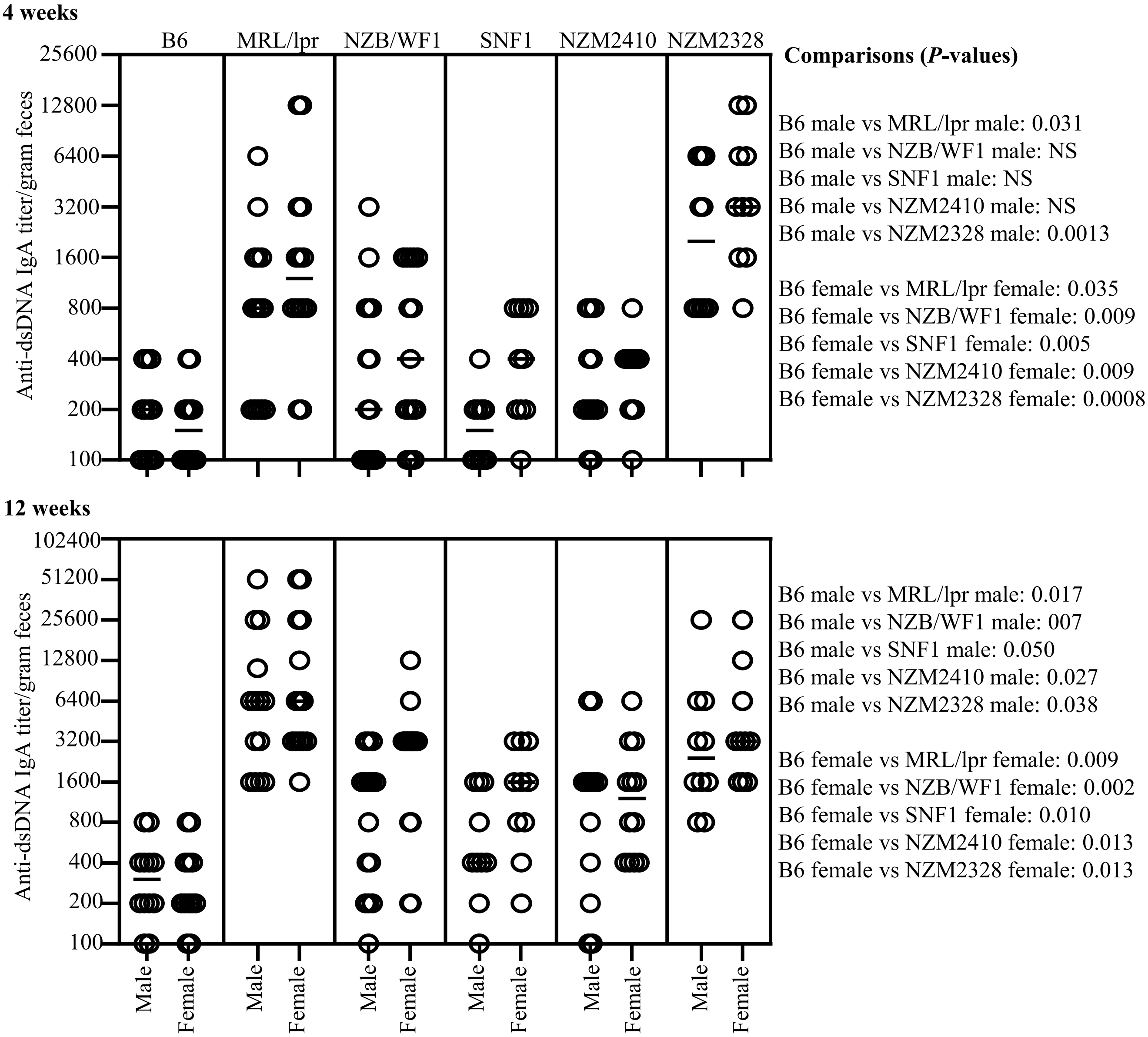


**Supplemental figure 3: dsDNA reactivity of IgA in the feces of lupus-susceptible and non-susceptible mouse strains.** Fecal samples were collected from individual mice of indicated strains and subjected to ELISA to determine dsDNA reactive IgA titers as detailed under the materials and methods section. dsDNA-reactive IgA per gram feces (collected at 4 and 12 weeks of age) are shown. n= 12 males and 12 females for B6, 14 males and 14 females for MRL/lpr, 15 males and 15 females for NZB/WF1, 10 males and 10 females for SNF1, 12 males and 12 females for NZM2410, and 10 males and 10 females for NZM2328 strains. *P*-value (by Mann-Whitney test) of various comparisons are shown on the right. Similar differences were obtained when nucleohistone reactive IgA titers were compared (not shown).
